# Monitoring changes in the Gene Ontology and their impact on genomic data analysis

**DOI:** 10.1101/320861

**Authors:** Matthew Jacobson, Adriana Estela Sedeño-Cortés, Paul Pavlidis

## Abstract

The Gene Ontology (GO) is one of the most widely used resources in molecular and cellular biology, largely through the use of “enrichment analysis”. To facilitate informed use of GO, we present GOTrack (https://gotrack.msl.ubc.ca), which provides access to historical records and trends in the Gene Ontology and GO annotations (GOA). GOTrack gives users access to gene- and term-level information on annotations for nine model organisms as well as an interactive tool that measures the stability of enrichment results over time for user-provided “hit lists” of genes. To document the effects of GO evolution on enrichment, we analyzed over 2500 published hit lists of human genes (most over 9 years old). 53% of hit lists were considered to yield significantly stable enrichment results. Because stability is far from assured for any individual hit list, GOTrack can lead to more informed and cautious application of GO to genomics research.

## Introduction

The Gene Ontology (GO) has been widely adopted by computational and experimental biologists and Gene Ontology annotation (GOA) of genes is one of the most prominent descriptive features of major genome databases. The original paper describing GO (Ashburner et al., 2000) is among the most cited papers in the biomedical literature (over 14,000 citations, Clarviate Analytics Web of Science, accessed 1/2018). The popularity of GO is in large part due to the challenge of interpreting data generated from high-throughput technologies such as gene expression profiling.

In a typical simple setting, researchers contrast a genome-wide feature (e.g., gene expression levels or genetic association) in two experimental conditions and generate a list of genes, either ranked across the whole genome, or in the form of a “hit list” of selected candidates. Another way such lists can be generated is by clustering, such as using protein interaction networks or coexpression; or by selecting genes harboring potentially pathogenic variants in cohort-based genome sequencing. To help extract biological meaning from those rankings and hit lists, it is now standard practice to use GO annotations in an “enrichment” framework. The widespread use of these methods motivates increasing the ability to understand their underpinnings.

Despite the importance of GO, many users likely have little understanding of how it was developed and how it changes over time, despite some effort on the part of the GO Consortium (GOC) to disseminate such information (Blake, 2013; Gaudet and Dessimoz, 2017; Huntley et al., 2014a). This is in part because there is no resource available to directly assess changes. Our goal in this paper is to fill this gap and provide some insight into the actual impact of changes on data analysis.

The structure, content and curation of GO/GOA is the essential backdrop for the work we present so we review it briefly. GO is organized into three sub-ontologies, representing Biological Processes, Molecular Functions and Cellular Components. Collectively these currently encompass over 47,000 concepts, arranged in a directed acyclic graph (like a tree, but with the potential for multiple paths from any leaf to the root). It is important to distinguish the GO itself from the annotations (GOA), which connect genes to terms in GO. Both GO and the annotations change over time as curation is performed.

Curation is managed through the Gene Ontology Consortium, in which member organizations such as model organism database curation teams provide annotations to a central repository. Genes may be associated with terms in the ontology using either manual curation (associated with a specific reference to the literature or based on a computational analysis reviewed by a curator) or “automatic” annotations that are not reviewed by curators. The different types of associations are represented by evidence codes, for example the automatic annotations receive the code “IEA” (“Inferred from electronic annotation”).

Annotations created by the curation process are referred to as “direct annotations” because they explicitly associate a GO term with a gene. Genes are also associated with terms indirectly via the graph structure of GO, referred to as inference. Thus, a gene that is directly annotated with the term “protein tyrosine kinase” is also implicitly annotated with the term “protein kinase” because that term is a parent term of “protein tyrosine kinase”. When the operation of propagating direct annotations through the GO hierarchies is completed (“transitive closure” in graph theory terminology), the number of annotations available greatly increases, albeit at a range of granularities. These “indirect annotations” (also referred to as “inferred” or “propagated”) are as valid as direct annotations because GO enforces a “true path” rule (Consortium, 2001). In most analyses, it is important to use propagated annotations (the combination of direct and inferred annotations) (Rhee et al., 2008).

Assessments of GO/GOA have recently turned to considerations of changes over time. For example, we quantified the effect that annotation have on the apparent (annotated) function of genes, showing that on average changes over short periods (months) are minor, but changes over longer periods are much more substantial (Gillis and Pavlidis, 2013). This and other work has shown that GO enrichment results may not be stable over time. However, the effects of changes are not likely to be uniform across data sets nor easily predictable. Indeed, previous studies have been either anecdotal (considering a single or just a few examples (Alam-Faruque et al., 2011; Clarke et al., 2013; Groß et al., 2012; Wadi et al., 2016)), with the largest study analyzing around 100 (Tomczak et al., 2018), or yielded mixed findings. Groß et al. (2012) found that enrichment results were stable based on analysis of two hit lists. Alam-Faruqe et al. considered changes in results to be improvements due to focused curation, based on analysis of two data sets. Others have emphasized instability (Tomczak et al., 2018; Wadi et al., 2016) or reported mixed impacts (Clarke et al., 2013). Given this variety of results and interpretations, there is clearly a need for researchers to assess the stability of their own specific enrichment results.

Here we report the development and application of a database (GOTrack, gotrack.msl.ubc.ca) that contains historical information on GO going back to the early 2000s for human and major model organisms. The GOTrack web site enables quick exploration of GO and GO annotations over time, and evaluation of how changes impact interpretation of analyses derived from GOA. Using the data in GOTrack, we present several analyses of trends in GO annotations, complementing earlier work. We performed a large-scale analysis of enrichment analysis results over time, using a large corpus of over 2500 “hit lists”. We confirm that GO enrichment analysis results can change over time. However, many were stable by objective measures even over time spans of greater than 10 years. It is our hope that GOTrack will enable more critical use of GO by biologists and computational researchers.

## Results

### Construction and overview of GOTrack

We used data representing ontologies and annotations for nine organisms, dating as far back as 2001. Annotation data were not available for all organisms for all dates, with complete data for all nine organisms from November 15 2011 onwards. In total the data encompasses 206 monthly versions of GO and 1545 species-specific monthly editions of GOA yielding a grand total of 206,894,446 GO annotations (as of January 2018). Our overall procedures are outlined in Figure 1 (see Methods and further information is available on the GOTrack web site). The resulting database is complex and rich, with extensive information available at the gene or GO term level. While the web interface is the most complete and detailed way to interact with the data, we also offer a RESTful API to enable programmatic access to the data. Via this API, users can download GO annotations for a taxon, as well as GO, for any selected point in time (https://gotrack.msl.ubc.ca/downloads.xhtml). GOTrack does not contain all information on GO/GOA and thus complements other resources such as QuickGO (Binns et al., 2009) and AmiGO (Carbon et al., 2009).

**Figure 1:**
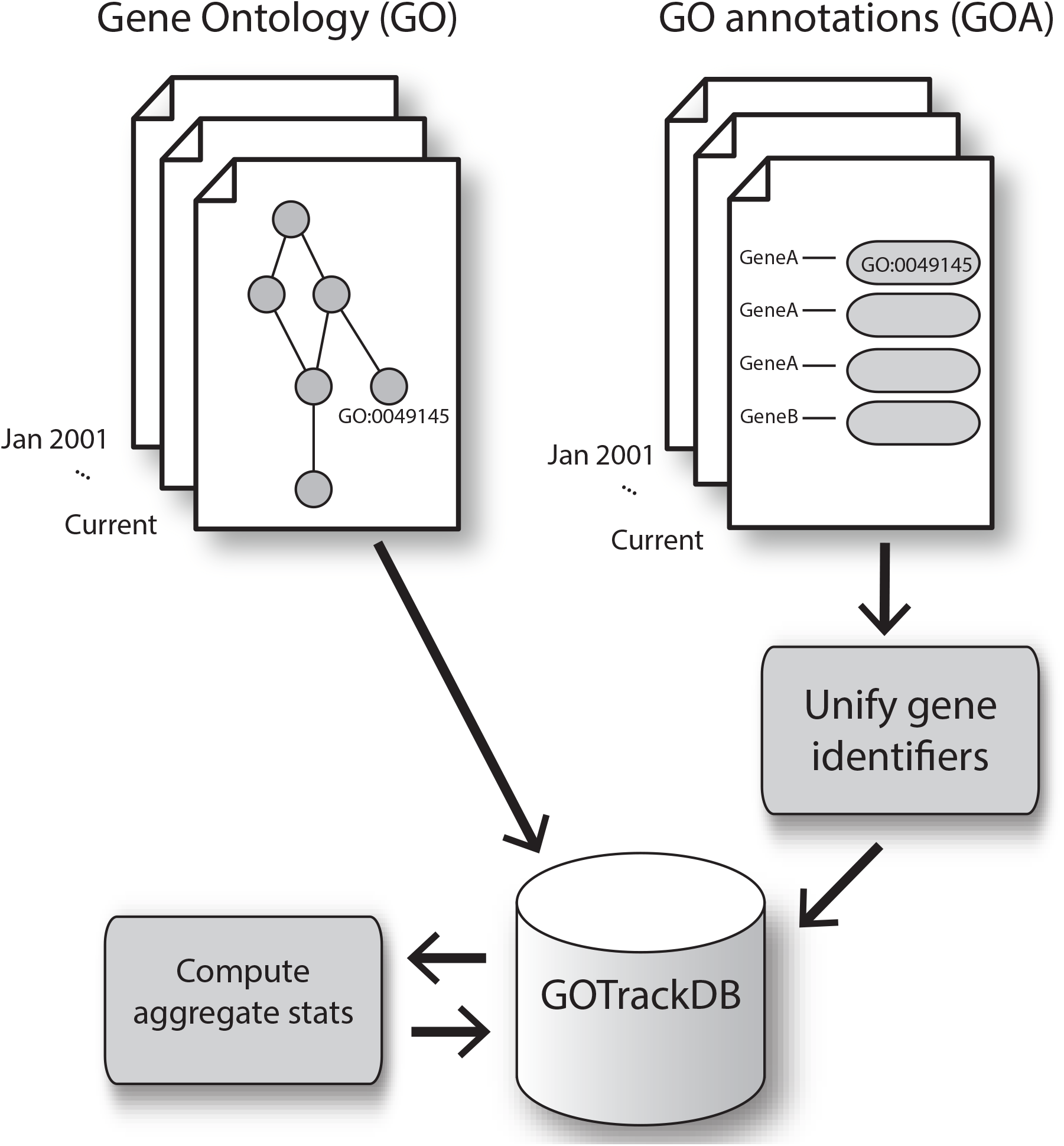
Overview of approach in constructing GOTrack. GO terms and GO annotations were obtained, matched by date, and harmonization of gene identifiers. Precomputed summary and aggregate statistics supplement the fine-grained information stored in the databases.

The GOTrack web interface offers views of history at the gene level, and at the GO term level. A third view provides a “global overview” of trends according to a variety of statistics. Finally, we offer a web tool to track changes in GO enrichment results over time. In this paper we provide only a high-level overview of basic functionality and readers are invited to explore the web interface for more information.

Figure 2A shows an example of one type of data offered in the gene view, for the human gene GRIN1 (glutamate ionotropic receptor NMDA type subunit 1; https://gotrack.msl.ubc.ca/genes.xhtml?accession=Q05586 and supplementary Figure 1A). The plot shows the number of GO terms directly annotated to the gene, with the mean of all genes from the same organism plotted for comparison. GRIN1 is consistently more highly annotated than the average, and its trajectory is typical in that annotations rise over time, interrupted by drops and recoveries. In general, such changes can be due to either annotation curation - addition or removal of terms annotated to the genes - or changes in the structure or content of the GO itself such as addition of terms or relations. The GOTrack interface also allows users to inspect changes in the use of evidence codes used to support an annotation, and directly compare annotations for a gene at up to four time points.

**Figure 2:**
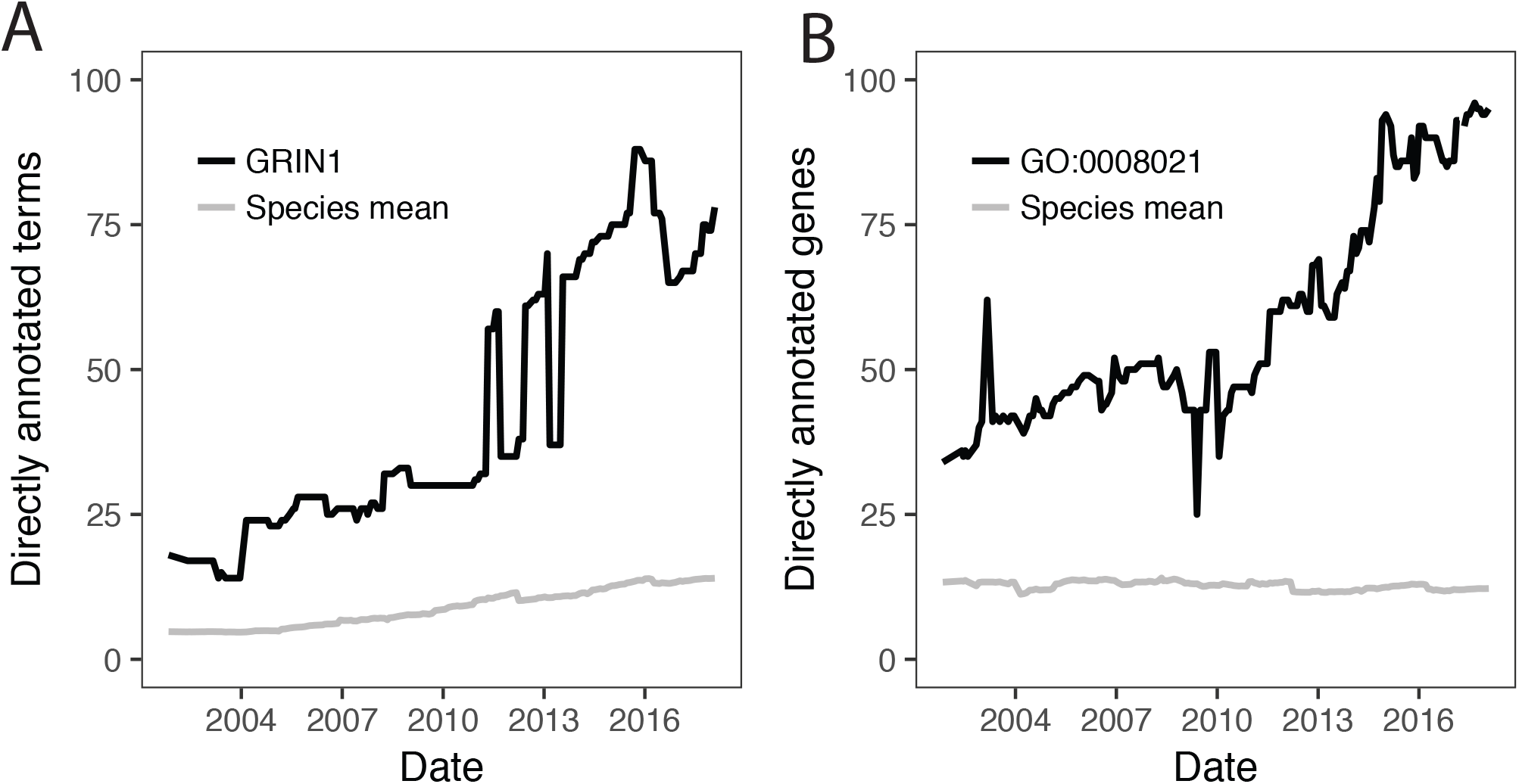
Examples of information provided by GOTrack for genes and terms. A. Number of terms directly annotated to the human gene GRIN1. Large drops and rises are observed superimposed over a general gradual increase in annotation since 2002 (black). In this example the large shifts are not accompanied by corresponding shifts in the species average (grey). B. Number of human genes directly annotated with the term “synaptic vesicle” (GÒ:0008021) over time, again showing transient drops and rises. Data from GOTrack was replotted for presentation. For corresponding screenshots, see Supplementary Figure 1.

To help users interpret the changes in number of terms over time, we provide additional plots of statistics derived from the annotations. The first of these is of multifunctionality (Ballouz et al., 2016; Gillis and Pavlidis, 2011), which is related to the number of terms annotated to a gene, with a weighting to account for term specificity (where specificity is defined by how many genes are annotated with the term; see Gillis and Pavlidis 2011 for details). This more precisely captures how heavily annotated the gene is relative to other genes. The second derived statistic is semantic similarity. As time passes, changes in annotations can cause a gene to change “functional identity” (Gillis and Pavlidis, 2013). To quantify this effect, we plot the Jaccard index between the annotations in the current edition to each previous edition. These and other plots and tables are presented on the web page for each gene.

The term-level view provides information on how a GO term has changed over time. This includes how many genes were annotated to it either in total (Figure 2B) or broken down by evidence type https://gotrack.msl.ubc.ca/terms.xhtml?query=GO%3A0008021 and Supplementary Figure 1B) as well as changes in the GO structure that impact the terms relationships. Finally, the Global Trends page (https://gotrack.msl.ubc.ca/trends.xhtml) shows species-level summaries of the numbers of annotated genes, genes annotated per term, annotations per gene, and the size of GO itself.

### Long-term trends in GOA

In this section we present some analysis of the data in GOTrack, focusing on annotations (rather than GO itself). As noted, genes vary strongly in how highly annotated they are, due to varying degrees of experimental and curation attention paid to the gene as well as potentially true biological differences in multifunctionality (Gillis and Pavlidis, 2011). We previously reported that this bias tends to persist, that is, genes which are relatively highly annotated tend to stay that way (Gillis and Pavlidis, 2013). We confirmed this is still the case five years later. For example, if we rank genes by how many direct annotations they have, the ranking at the earliest time point is correlated with the ranking at the latest time point (human: Spearman rank correlation 0.52; mouse 0.43; Arabidopsis 0.53). Thus we confirm that genes are not just unequal in their annotation, but that this inequality is stable over long periods.

The jumps seen in individual genes (e.g. Figure 2A) are not all independent events, as the course of the species-wide averages also has discontinuities (Figure 2A, grey). This is also apparent in a principal components analysis of the direct count matrix (Supplementary results, Supplementary Figure 2) We investigated this more completely in all nine GO Track organisms at the level of total gene coverage (Figure 3A), genes annotated per term (Figure 3B), direct annotations per gene (Figure 3C) and inferred annotations per gene (Figure 3D). This reveals that large jumps and drops are sometimes simultaneously observed in multiple, or even all species. One such notable event was a rapid increase in the number of annotated genes starting March 2011 for Arabidopsis, mouse and zebrafish (Figure 3A). Inspection of update reports from GOA (https://www.ebi.ac.uk/GOA/news)leads us to speculate that the jump might be due to the Reference Genome Annotation Project (The Reference Genome Group of the Gene Ontology Consortium, 2009). Another dramatic event was a large drop in the mean number of direct annotations per gene in March 2012 for all species (Figure 3C). The jump is not visible in the plots for indirect annotations (Figure 3D). This would be consistent with a large-scale purging of redundant annotations (rejecting higher-level terms that are inferable from more specific terms). Other jumps are species-specific, such as the large increase in Arabidopsis genes annotated per term in early 2012, followed by a large drop in late 2015 (Figure 3B).

**Figure 3:**
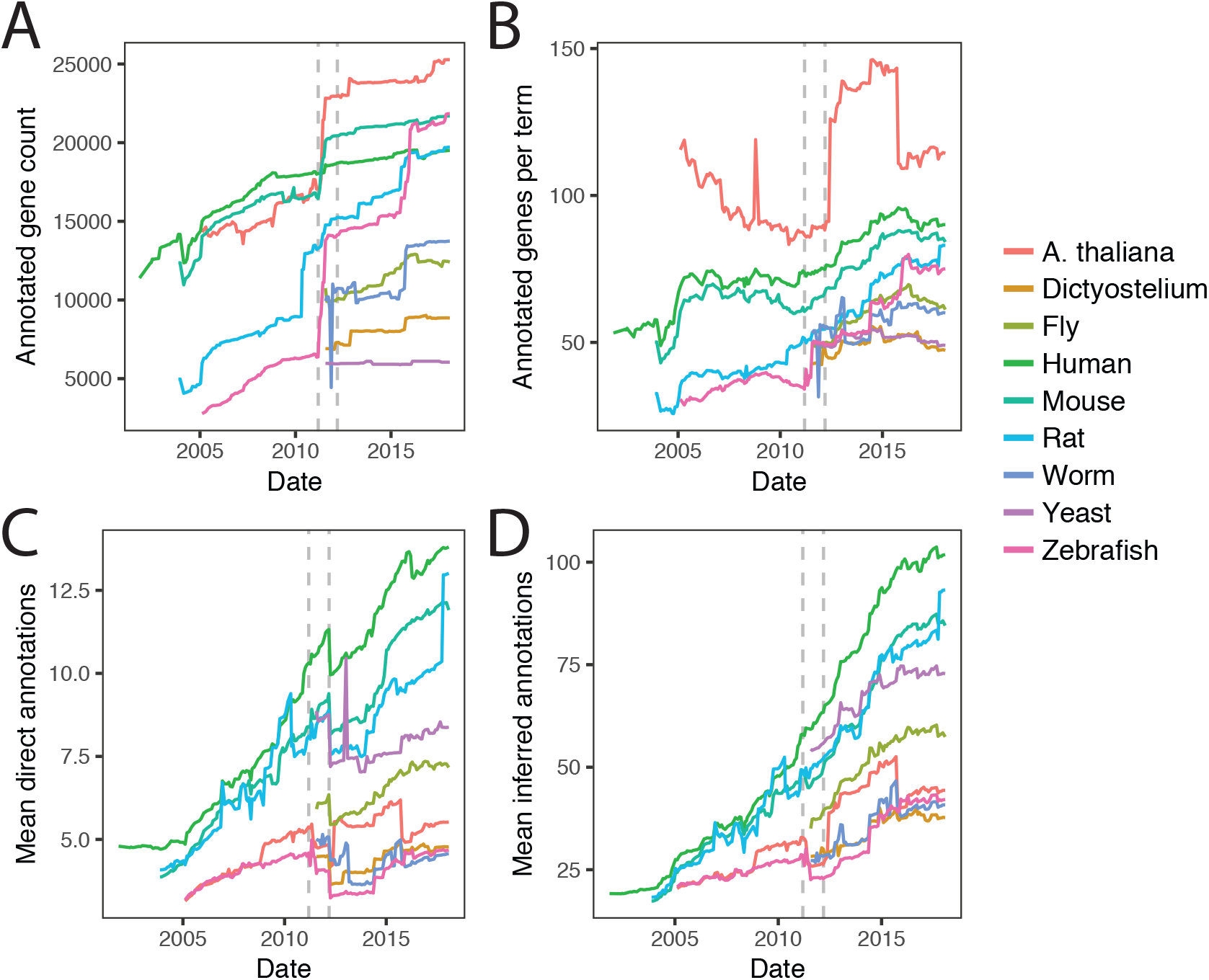
Trends in taxon-wide annotation statistics. A: Number of annotated genes. B: Mean annotations per term (inferred + direct). C: Mean number of direct annotations per gene. D: Mean number of inferred (including direct) annotations per gene. Times of prominent discontinuities affecting multiple species in A and C are marked by dashed grey lines in all four panels.

At the gene level, large shifts in the numbers of annotations can be due to removal and replacement of annotations for the same term - a phenomenon we call “annotation churn”. For example, for the human gene ACTC1 (https://gotrack.msl.ubc.ca/genes.xhtml?accession=P68032), there is a pronounced rise in annotations in mid-2007, with a one-month dip in May 2008 (see screenshots in Supplementary Figure 3). GOTrack makes it easy to drill down into details. By examining the tabular results (Supplementary Figure 3A), it is found that one of the terms that was deleted during the dip was “apoptosis” (GO:0006915). Viewing the annotation history for that term on the gene, we see that the term was repeatedly added and removed (in 2007-2008), with the evidence code “IEA”. In June 2008 the term was annotated to the gene with a higher-grade curator-reviewed evidence code (ISS), where it remained (the term was also renamed to “apoptotic process”) - until it was removed again in December 2017 (Supplementary Figure 3B).

### Tracking enrichment results

In addition to the exploratory aspects described so far, the other major component of the GOTrack system is an analysis tool which performs enrichment analysis at multiple time points (https://gotrack.msl.ubc.ca/enrichment.xhtml; Supplementary Figure 4). The key idea is to observe whether an enrichment result is stable relative to a given point in time. The main input provided by the user is a “hit list” of genes. The output includes plots and detailed tables to help interpret the results and judge whether the results change over time. This includes direct comparisons of “before and after” sets of enriched terms. The measures we use for this comparison are discussed in the next section and in Methods. In addition to these statistics that summarize the overall stability of the results, the web interface provides term-level stability measures. This makes it easy to see whether a term has been consistently “significant” over past editions.

The enrichment tool has some limitations: we use a simple over-representation method (as do many tools including the popular DAVID (Huang et al., 2009), and the “background” set of genes is not settable by the user: it is the set of all genes annotated at the particular time point. But because GOTrack provides downloads of GO and GOA for any date, users can confirm findings with software of their choice, provided it allows user-provided GO and GOA as inputs (such as ErmineJ, (Ballouz et al., 2016), whose annotation input format is directly supported).

### Evaluating the stability of enrichment results

We hypothesized that changes in GO/GOA over time could cause changes in enrichment results to such an extent that they would be effectively unrecognizable and lead to a different interpretation of the results; as described in the introduction, previous studies of this question yielded somewhat mixed results on small numbers of test hit lists. In our approach to this question we used a corpus of gene lists from the Molecular Signatures Database (MSigDB) (Subramanian et al., 2005). These are divided into two groups (after filtering, see Methods): 1327 curated “canonical pathways” (CP) and 2573 “chemical and genetic perturbations” (CGP). The latter correspond to published hit lists of the type usually investigated with enrichment analysis. We took advantage of the fact that each CGP hit list is associated with a publication, allowing the opportunity to see if the enrichment results obtained around the time of publication would have changed in the interim. We predicted the CP lists of established pathways would be more stable compared to the experimental CGP hit lists. The limitation of the MSigDB corpus is most of the publications are not very recent (median 11 years; range 0.4-16, 90% are >9.2 years old) and we have done little investigation of short-term stability.

For each hit list or pathway, we compute results of an enrichment analysis as it would have appeared at the GO/GOA edition nearest to the source publication date (see Methods for details). We then repeated the enrichment analysis using the most current GO/GOA edition (January 2018). This results in a range of timespans to have passed following publication. For the CP set, which do not all have an associated date, we computed results for the most recent GO/GOA edition and the earliest date available (January 2001). We used this extreme date for comparison because we expected the CP set to have greater stability, so comparing to the earliest date is the “worst case scenario” for comparing to the experimentally-derived CGP sets.

Our first key observation is that on average for the CGP hit lists, the number of significant terms goes up dramatically (from 21 ±32 terms to 110±136 terms, mean ± standard deviation; p<10^−15^, Wilcoxon rank sum test). The values are highly correlated (Figure 4A): hit lists that had few significant terms at the time of publication (henceforth t_0_) had relatively few at the most recent timepoint (t_now_) (rank correlation 0.54). These results also held for the Canonical Pathways (growing from 37±59 to 246±216 terms, correlation 0.57). It is likely that these increases are not just due to increased annotation, but the growth of GO to over 47,000 terms of increasing granularity.

**Figure 4:**
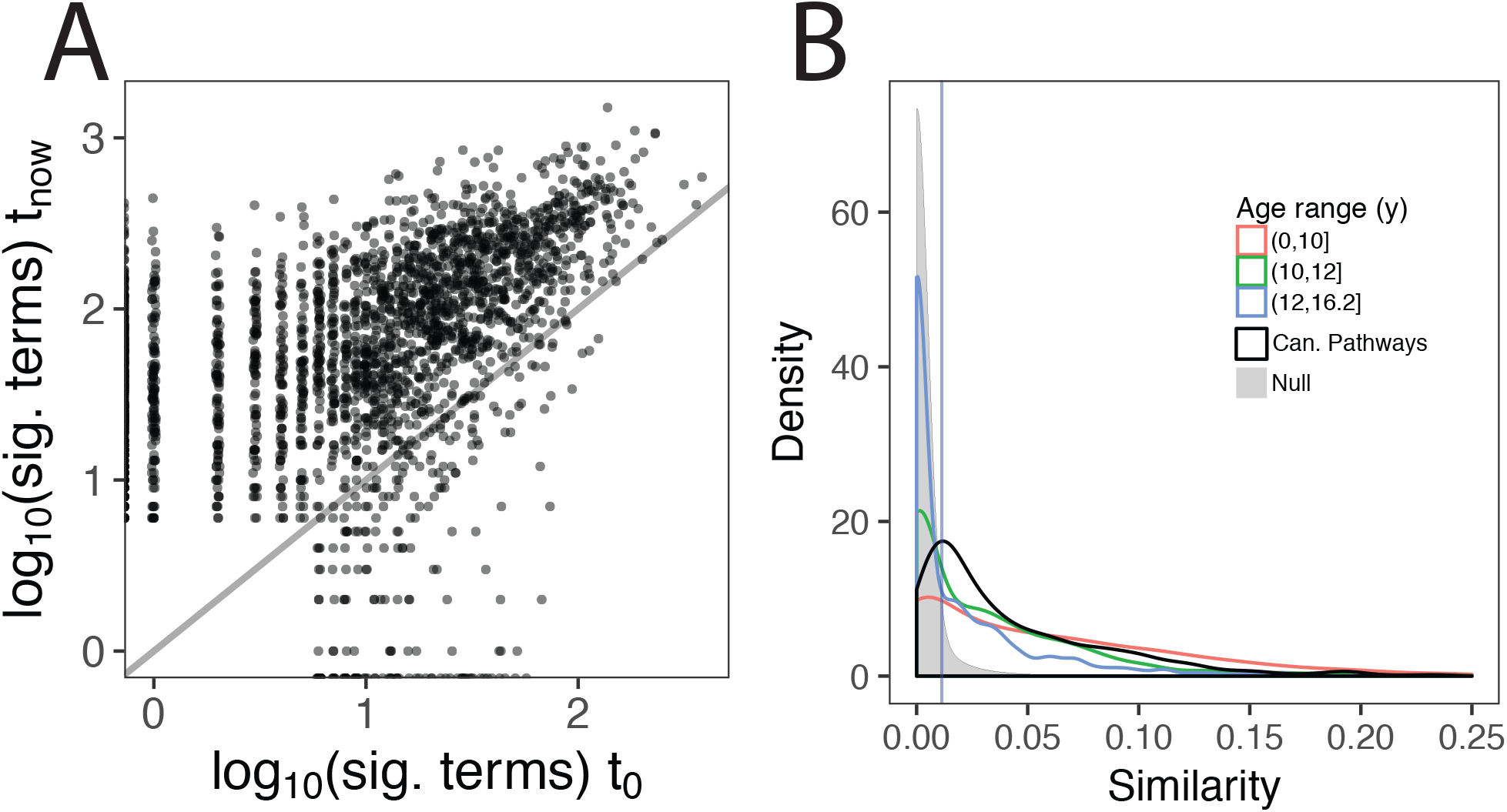
Stability analysis of 2573 published hit lists A. Change in number of significant GO terms. Each point is one CGP hit list. Points are jittered to reduce overplotting. B. Similarity of enrichment results, using the complete Jaccard index. The CGP hit lists are binned into most recent (orange), old (green) and oldest (blue). The distribution for the canonical pathways is in black. The blue vertical line indicates the 95°ile of the null.

The explosion in the number of significant terms is an obvious form of instability, but of course what matters more is whether the enriched terms resemble each other at t_now_ compared to t_0_. To evaluate this, we did direct comparisons of the enriched terms associated with each hit list (at t_0_ and t_now_), using the Jaccard index (see Methods and Supplement). The Jaccard index was calibrated using a null distribution created by comparing pairs of unrelated hit lists (see Methods). To simplify the analysis, we binned the CGP hit lists by age into three groups of similar numbers of hit lists: up to 10 years, 10-12 years, and 12-16 years.

The results are shown in Figure 4B. Overall, 53% of the CGP hit lists had results which were more similar than 95% of the null trials. This fraction is much higher for relatively recent lists (71%, N=640) and lower for the older lists (55% for the middle tranche, N=960; and 38% for the oldest, N=973; Figure 4B). In comparison 75% of the Canonical Pathways remained above this threshold, despite most of the comparisons being done to the earliest possible time point. The overall rank correlation (unbinned) between stability and age is −0.34 (CGP; −0.39 for Canonical Pathways). This demonstrates that it is possible for results to maintain a substantial degree of similarity over periods of greater than 15 years, but that in general, drift in the semantic content of enrichment results is very substantial after 12-16 years and is substantial but less striking at shorter time spans (<10 years). In the Supplement we present examples of hit lists yielding high or low stability (Supplementary Results and Supplementary files).

A notable feature of the data shown in Figure 4B is that very low values of the complete Jaccard index were statistically significant. This shows the importance of using a null distribution to calibrate the scores, but clearly leaves something to be desired as a Jaccard index of 0.01 seems negligible. However, this effect is due in large part to the increase in the number of terms over time (Figure 4A), guaranteeing that the Jaccard index will drop. In attempts to explore this further, we tested six variants on the Jaccard index (see Supplement). While some of the alternatives have scales that are more intuitively matching expectations of what “stable” would represent on a scale of 0-1 (e.g., with 95%ile of the null equal to 0.41), the findings are qualitatively similar to the complete Jaccard (data for two additional measures are shown in Supplementary figure 5). Several of these alternative measures are implemented on the GOTrack web site. These measures are discussed further in the Supplement in the context of examples, along with discussion of the subjective nature of comparing enrichment results in an exploratory analysis.

We looked for factors that might contribute to stability. For the CGP hitlists, the number of genes in a hit list was not strongly predictive of Jaccard stability (rank correlation 0.18). It was only modestly correlated with the mean number of directly annotated terms (-0.12) or mean multifunctionality of the genes in the hit list (-0.09). There were more obvious trends for the canonical pathways lists, which have higher stability than the CGP lists on average, despite the (artificially) long time passed between t_0_ and tnow (over 12 years; Figure 4B). The number of direct annotations per CP is higher (36 vs. 25.4 for CGP). However, this does not appear to explain the overall higher stability of the CP lists, because the we get the same result for the subset of CP that has <35 mean direct annotations (mean of 22.9; correlation is −0.48; overall correlation −0.46). Thus hit lists that have more highly annotated genes have a tendency to be less stable. But given these low correlations (-0.12 for the CGP set) and without further insight, it appears to be difficult to predict (even in hindsight) which hit lists will yield stable results.

## Discussion

In this work we present GOTrack, which to our knowledge is the only resource available that allows easy access to historical data on GO/GOA, and the only that allows inspection of the effects of changes over time on enrichment result stability. Our analyses further highlight the necessity for users of GO/GOA to be cautious in their interpretation of any GO annotation, and to temper whatever trust they have in GO enrichment results.

Our evaluation of the stability of enrichment results differs in several important ways from earlier efforts. First, we matched GO and GOA for each time point (rather than fixing either GO or GOA while varying the other), which we feel is more realistic. We also analyzed a much larger number of hit lists (>2500 vs. a maximum of ~100 (Tomczak et al., 2018)) and considered time of publication to ensure comparisons were also realistic. But perhaps most importantly, we used a null distribution to calibrate the similarity measures, providing improved objective measures of what qualifies as stability. Overall our results are more optimistic about stability than Tomczak et al. (2018). Regardless, we concur with previous reports that changes in GO/GOA can make a substantial difference in results, but because of the high degree of variability and difficulty in finding fully satisfying quantitative measures that are often interpreted subjectively (see Supplement for discussion), our recommendation is that users of GO should judge for themselves by using GOTrack. Researchers who are reporting enrichment analyses can check which terms have been stable (for example, over the last five years). This provides a principled way to help narrow down complex enrichment results, a problem that many users of enrichment analyses struggle with.

An obvious limitation is GOTrack cannot see into the future. While the stability of any particular GO enrichment result might be high or low when looking backwards in time, it is generally impossible to know whether it will remain to be stable because knowledge of biology as represented in GO/GOA is a work in progress. Indeed, we found it is difficult to predict which hit list will give stable results. The strongest clue we could identify is how well annotated the genes in the hit list are: hit lists with highly annotated genes (mean direct annotation count) tend to be less stable. We speculate that this is because highly annotated genes have more changes to their annotations, which can drive shifts in enrichment results, but we have yet to explore this further and in any case the relationship is not strong enough to be usefully predictive. In addition we did not assess other possible factors influencing stability such as evidence codes (Yu et al., 2017), a topic we leave for future research.

GOTrack has some limitations that may be addressed in the future. The enrichment tool uses a simple method and does not implement algorithms to asses multifunctionality biases (Ballouz et al., 2016). Our data on GO/GOA is not complete: we did not import all of the fields from GO annotations files, the most useful of which for our purposes might be the annotation source. However, the granularity of source annotation is limited. Notably, annotations coming from the Reference Genome Project (The Reference Genome Group of the Gene Ontology Consortium, 2009) are not identified so we were unable to establish any specific impact this may have had on the events of early 2012 (Figure 2C). Finally, the recently added concept of annotation extensions (Huntley et al., 2014b), which provide context for an annotation (for example, a cell type) are not handled by GOTrack.

The evolving and incomplete nature of GO/GOA has always been inherent and is well understood by the GO community. But it is seemingly less appreciated more broadly. For example, the extremely popular enrichment tool DAVID (over 32000 citations as of May 2018, https://david.ncifcrf.gov/) did not update its GO annotations for nearly seven years, an eon in GO history (and at this writing DAVID has not been updated for nearly two years [https://david.ncifcrf.gov/content.jsp?file=release.html]). We find it interesting that there wasn’t a massive outcry in response to the use of such out-of-date GO annotations, suggesting either ignorance or apathy. While it might seem obvious that one would always want to use the latest GO annotations, this can be questioned. GO/GOA can change dramatically in a see-saw fashion over a period of months, suggesting that not all changes are improvements. Furthermore, we report a strong tendency for hit lists to yield ever more significant terms over time (Figure 4A), and it is not clear this comes with any increase in useful information. It could be that using GO/GOA from an earlier, simpler era might be beneficial for enrichment analyses (using a GO slim [http://www.geneontology.org/page/go-slim-and-subset-guide] may approximate this concept). While we may not be able to settle that question here, it is clear that whatever version of GO/GOA is used, it cannot be treated as a gold standard. Enrichment analysis should be considered exploratory, and never used as a primary finding (Sedeño-Cortés and Pavlidis, 2014). Computational researchers should also be cautious in using GO/GOA as an optimization target when developing and evaluating algorithms, especially since changes over time are not the only concern (Gillis and Pavlidis, 2011, 2013).

GOTrack should be a valuable resource for biologists to gain a greater understanding of where GO annotations come from and how they change over time, as well as their impact on the major use case for GO/GOA, enrichment analysis. Our analysis of the data in GOTrack also revealed a number of interesting features, and it is likely that deeper analyses can be used to gain more insight into patterns of curation that might influence future efforts.

## Materials and Methods

### Gene Ontology

Historical Gene Ontology files were retrieved from ftp://ftp.geneontology.org/go/ontology-archive/, specifically: Dates between 2001-01-01 - 2004-03-01 were obtained from separate process.ontology.<date>.gz, function.ontology.<date>.gz, and component.ontology.<date>.gz files and subsequently combined. Dates between 2004-04-01 and 2006-10-01 were obtained from gene_ontology.obo.<date>.gz. Dates after 2006-10-01 were obtained from gene_ontology_edit.obo.<date>.gz. These files exclude relationships that cross the three GO aspects and we restrict our analysis to IS_A and PART_OF relationships only.

### Gene Ontology annotations

Historical species-specific annotation files were retrieved from ftp://ftp.ebi.ac.uk/pub/databases/GO/goa/old/<species>/, specifically: Dates between 2001-11-02 and 2016-05-09 were obtained from gene_association.goa_<species>.<edition>.gz. Dates after 2016-05-09 were obtained from a combination of goa_<species>.gpi.<edition>.gz and goa_<species>.gpa.<edition>.gz files. Mapping of historical annotations to a release of the Gene Ontology was done by selecting the ontology with the closest release date before that of the annotation file. Annotations were propagated up the GO graph as per the “true path rule” (Consortium, 2001). To convert release editions to dates, prior to edition 135 (July 2014) the release number of the file is compared to the dates given on the GOA news site (https://www.ebi.ac.uk/GOA/news). For edition 135 onwards we use the date provided in the files. We note that there are some gaps in the available data, especially at early time points. For example, we lack data for human for September and October 2002. In addition, the spacing of dates is not uniform; while the median inter-edition gap is 28 days, there are a few gaps that are smaller (minimum 13 days) or correspondingly larger (e.g. 40 days).

### Mapping of gene identifiers over time

Gene product annotations are tracked historically using their associated UniProt accession number(s) (Bateman et al., 2017). Each gene product in UniProt has a unique primary accession, called the ‘Primary (citable) accession number’. In addition to this, a gene product may also have secondary accession numbers which could have been created historically from merges and/or splits. During a merge, the first accession is retained as the primary while all others become secondary. During a split, a new primary accession is created for all products involved while their original accessions are retained as secondary. An accession is only deleted when its corresponding entry has been removed from UniProt. The mapping of primary to secondary accessions is retrieved from ftp://ftp.uniprot.org/pub/databases/uniprot/knowledgebase/docs/sec_ac.txt. This mapping allows us to find the current primary accession of a historical annotation.

### Enrichment analysis

GOTrack implements over-representation analysis using the hypergeometric distribution (Ballouz et al., 2016). The background is the set of all annotated genes (for the time point being analyzed). For analyses presented in the paper, terms with between 20 and 200 genes were included, and only Biological Process terms were considered. The false discovery rate was controlled at 5% using the method of Benjamini and Hochberg (Benjamini and Hochberg, 1995). The GOTrack enrichment tool allows these parameters to be varied by the user.

### Data analysis

Many of the analyses described are based on files available via https://gotrack.msl.ubc.ca/downloads.xhtml including the “summary” files by edition, terms and genes. Analyses were conducted with custom scripts written in R (Team, 2016; Wickham, 2009) and python. Code and all data files needed to reproduce the analyses are provided at http://hdl.handle.net/11272/10596. Correlations are Spearman Rank correlations except where indicated otherwise.

### Analysis of MSigDB hit lists

The MSigDB C2 collection (Subramanian et al., 2005) was downloaded from http://www.broadinstitute.org/gsea/msigdb/genesets.jsp. This corpus is divided into a set of “canonical pathways” (CP) and “chemical and genetic perturbations” (CGP). For the CGP hit lists, the publication associated with each hit list was extracted, and the date of publication (t_0_) was used to identify the nearest matching version of GO/GOA in GOTrack. Each hit list was analyzed for enrichment as described above, for t0 and a recent comparison time point (January 2018, t_now_). We analyzed 2573 CGP hit lists that yielded at least five significant terms at either (or both) t_0_ and the comparison time point. CP lists (n=1327 after filtering) were treated the same way, except t_0_ was fixed at Nov 21 2005 (the mean date for the CGP lists).

To compare two sets of enrichment results, we explored several measures (see Supplement) but focus on a standard Jaccard index: |*E*0 ∩ *E*1|/|*E*0 ∪ *E*1|, where E0 and E1 are the sets of all significantly enriched GO terms for the same input hit list at two time points (“complete Jaccard”). The primary alternative measure we examined was a modified Jaccard that examines only the top five terms plus their inferred parent terms (“top-term-parents Jaccard”), similar to the measure proposed by (Mistry and Pavlidis, 2008). See the supplement for details and discussion.

To generate a null distribution, we compare enrichment results from pairs of randomly-selected hits lists (i.e., coming from different publications). Instead of comparing a hit list’s results for t_0_ to t_now_, the data are permuted so t_0_ of one hit list is compared to t_now_ for a randomly-selected hit list (with the same constraint that at least one of them must have 5 or more significant GO terms). We analyzed 1000 such permutations of the data and pooled them to generate the null distribution. This is an appropriate null because if two enrichment results from the same experiment (at two different time points) are less similar than what would be expected for two randomly picked independent experiments, we can say that the enrichment results are no longer similar according to the measure. This null also inherently addresses the tendency of some GO terms to recur more frequently than others in independent enrichment analyses (Ballouz et al., 2016).

### Implementation and availability

GOTrack is implemented in Java and JavaScript, and uses the PrimeFaces framework, with a MySQL database. The open source Highcharts (highcharts.com) visualization library is used for plotting. GOTrack is open source software (https://github.com/PavlidisLab/gotrack) released under the Apache 2.0 license. The data in GOTrack are automatically updated monthly. Because of the lag in when data are available from GOC, data for a given date appears in GOTrack up to 2 months after the stamped date.

## Contributions

PP conceived of the project and provided oversight. AES-C and PP developed the original GOTrack web site concept. MJ implemented GOTrack based on a prototype developed by AES-C. PP, MJ and AES-C performed analyses and drafted the manuscript.

## Acknowledgements

We thank the GO Consortium for their efforts in creating GO/GOA. We are also dependent on the work of UniProt for protein identifiers. We thank Pascale Gaudet and Jesse Gillis for discussion, members of the Pavlidis lab for discussion and comments on the draft manuscript, and Dmitry Vavilov for technical support.

## Supplementary files

Supplementary results and discussion

Supplementary Figure 1: Screen shots of the gene and term views in GOTrack Supplementary

Figure 2: Principal components analysis of the direct annotation count matrix Supplementary

Figure 3: Screen shots showing annotation history tracking for a gene (annotation churn) Supplementary

Figure 4: Enrichment web interface

Supplementary Figure 5: Analysis of MSigDB lists using alternative similarity measures Supplementary

Figure 6: Correlations of stability measures

Supplementary Files 1-3 (APPEL_IMATINIB_RESPONSE.enrichment.xlsx, BENPORATH_ES_2.enrichment.xlsx, ONDER_CDH1_SIGNALING_VIA_CTNNB1.enrichment.xlsx): Examples of CGP enrichment results discussed in the supplement.

## Supplementary results and discussion for “Monitoring changes in the Gene Ontology and their impact on genomic data analysis”

### Principal components analysis (PCA) of the direct annotation counts

We performed a PCA on the direct annotation count matrix for the three species with the longest records in GOTrack (Arabidopsis, mouse and human). This matrix contains, for each gene at each time point (at monthly resolution), the number of directly annotated terms. Prior to PCA the data were scaled to force each gene to have the same mean (0) and variance (1), which eliminates the effect of different amounts of annotations overall per gene.

As expected the first PC (50-60% of the variance in the scaled PCA; >90% if data are unscaled) captures a general upward trend with some notable discontinuities (discussed in the main paper). Unexpectedly, other components in all three organisms had an oscillatory character with periods ranging from ~10 years (PC2) to ~2 years (e.g. PC 7 and higher) (Supplementary Figure 2, top). While we have not found a specific explanation for these patterns, our interpretation is that there is a subtle periodic character to curation efforts, including some short-term relative stability on the span of ~1 year interrupted by relatively large changes. This is also readily visualized in the direct annotation count correlation matrices (Supplementary Figure 2, bottom).

### Evaluating measures of stability

One challenge we encountered was finding a satisfactory metric for similarity of enrichment results that captures the way enrichment results are often interpreted, which is to say, loosely. Earlier work considered tended to focus on the exact terms which were “significant”. However, we found that it was common for results to have low similarities at the exact term level (e.g. Jaccard similarity of significant terms) while yielding a similar “biology impression”, especially for the top terms. To give a hypothetical example, it is unlikely a user would (or should) care whether the top enriched term is “BMP signaling pathway” or “cellular response to BMP stimulus” - the subjective impression is the same (we use the term “impression” in this manner throughout). For this reason, expecting that all significant terms match exactly might give an overly pessimistic view of the stability of the results, especially in light of our results showing that the number of enriched terms tends to increase over time. This phenomenon effectively guarantees that the Jaccard index will tend to drop, as novel terms are called significant in addition to previously significant terms. Relatedly, users often focus on top-ranked terms. Another factor is it is possible in theory for a term to be enriched at two time points, but the support for the enrichment comes from different genes. These considerations motivated us to implement seven similarity measures and to evaluate their properties in our analysis of the MSigDB gene lists. Because there is no gold standard, our evaluation is largely descriptive and somewhat subjective.

Our evaluations of stability are based on comparing two sets of GO terms, each of which is selected from an enrichment analysis, either by mere ranking (e.g. top five terms) or at an applied false discovery rate threshold (“significant terms”). All of the measures range from 0 to 1, with 1 meaning highest similarity. The baseline measure for comparing two sets is the Jaccard index, computed as |*E*0 ∩ *E*|/|*E*0 ∪ *E*|, where E0 and E1 are the sets of all significantly enriched GO terms for the same input hit list at two time points (“complete Jaccard”). This corresponds to the approach taken in (Tomczak et al., 2018). The other measures we considered are variations where we change the sets that are being compared. The variants are “Top terms Jaccard”, which is the Jaccard index of the top 5 terms of each ranking (or fewer, if there were less than 5 significant terms). “top term parents Jaccard” (“top parents” for short) is the same as “Top 5 terms” but expanded to include the ancestor terms in the GO hierarchy, similar to the measure proposed by (Mistry and Pavlidis, 2008). “Top gene Jaccard” compares the genes from the hit list annotated with the top 5 terms, rather than comparing the terms. This allows for the possibility that two editions could both have the same term called significant, but due to different annotated genes. The gene-based measure was intended as an adjunct to the others, rather than as a likely primary measure of stability. In addition, for each of the three methods we computed an asymmetric Jaccard (Tversky index):|*E*0 ∩ *E*|/(|*E*0 ∩ *E*| + |*E*0 − *E*1|), where E0 represents the earlier time point. If all the terms in E0 are in E1 (|*E*0 − *E*1| = 0) the denominator is maximized. The Tversky index focuses on how well the original set E0 was preserved in E1, so new terms that appear in E1 do not detract from the score (Tversky, 1977), which is potentially appropriate for the task of considering how well preserved results from a past date are now. In total we had seven measures, of which the four Jaccard-based are currently shown on the GOTrack web site. The web site also allows users to pick the time point to be used as a reference.

All of the measures we selected were at least moderately correlated (rank correlations for the CGP hit lists between 0.43 and 0.92, Supplementary Figure 6). We generated a null distribution as described in Methods for the complete Jaccard” index as well as for the “top-parents Jaccard” and the asymmetric “top-parents Tversky” and examined them in more detail. All three measures yield comparable results in the CGP hit lists (Figure 4B and Supplementary Figure 5). As reported in the main text for the complete Jaccard index, 53% of CGP hit lists exceeded the of 95%of the null trials. The value was 36% for top-parents Jaccard and 33% for top-parents Tversky.

We found the “top gene” methods (genes supporting the enriched terms) is useful as an adjunct to the other measures as a way to inspect stability at the level of genes after establishing stability of a term, as illustrated with the cases described above. But we do not recommend it as a primary measure of stability. Finally, the top-terms (without parents) methods were most highly correlated with the complete Jaccard (all significant terms), but also noisy in terms of variance over short time spans (low autocorrelation).

### Examples of stable and unstable enrichment results

In this section we describe some examples that illustrate our findings and highlight some of the limitations of the semantic similarity measures.

To give an example of a stable hit list, the CGP list “APPEL_IMATINIB_RESPONSE” (Appel et al., 2005), which dates to 2005 (33 genes), had results with a complete Jaccard similarity between t_0_ and t_now_ of 0.046, which appears negligible and gives a strong impression that the results have completely changed. However, this is well above the 95%ile of the null (0.011). For the top-parents Jaccard, the similarity is much higher (0.33, also well above the 95%ile of the null). Thus, objectively this hit list yields very significantly similar results in 2018 compared to those it would have yielded in 2005, but it is concerning that the values are low in an absolute sense. In situations like this where there is “statistically significant” stability relative to the null, the low absolute scores raise the question of whether the results are meaningfully stable for practical application. This brings us back to the point about whether results give the same overall impression, regardless of what the numeric scores reveal. By definition this is a subjective discussion and we have only done informal investigation.

Continuing with the example of APPEL_IMATINIB_RESPONSE, we find that 2 of the 9 terms which would be considered significant in 2005 were still ranked in the top 9 terms in 2018 (GO:0006664, GO:0006665), and altogether 5/9 are still significant. Of the other four terms, one was ranked 140 and no longer significant. Three (GO:0006869, GO:0044275, GO:0006643) dropped out of the analysis before the present, of which two would have been considered significant at the date of their last inclusion (dropouts occur if the gene set size goes outside the limit of 20-200 we set for the analysis). The reason the complete Jaccard similarity is so low (0.046) is there are now 104 significant terms rather than just 9. The reader should judge for themselves if the results are similar (Supplementary File 1), but this example is a good illustration of what we mean by “impression” - the similarity of the top of the ranking is very high, and many of the added terms are simply more specific variations on the themes captured by the top terms. Instead of just “membrane lipid metabolism” and “lipid transport”, the new list includes “sphingolipid catabolic process”, “phospholipid transport”, “fatty acid transport”, “positive regulation of lipid localization”, “lipid homeostasis” and so on. There is still a balance of completely novel-sounding terms such as “viral life cycle”, “lung morphogenesis”, “locomotory behavior”, “animal organ regeneration”, “negative regulation of MAP kinase activity” and “acute-phase response”, but these have no obvious coherence (in terms of overall impressions, and bearing in mind all these terms are picked up by the same 33-gene hit list from a study of monocytes). In agreement with these observations, the “top genes” Jaccard measure for this hit list is 0.73: in large part the same specific genes within the hit list are driving the enrichment observed. We made similar observations of many hit lists that have low numeric stability but give results that give a subjective impression of reasonably high stability.

The other side of this situation is whether objectively low scores (compared to the null) match subjective impressions of “instability”, as well. The answer is yes, but arguably less convincingly. For example, the hit list BENPORATH_ES_2 (Ben-Porath et al., 2008, 40 genes) has a complete Jaccard similarity between t_0_ and t_now_ of 0.0. At t_0_, the enriched terms included “DNA replication”, “mitosis”, “methylation”, and “epigenetic regulation of gene expression”. While none of these terms are enriched at t_now_, highly related terms such as “DNA replication initiation”, “mitotic nuclear division” and “gene silencing” are enriched (Supplementary File 2). In our hands, the top-parents Jaccard measure was more in line with subjective impressions of a fair degree of similarity, with a significantly high value of 0.27 in this case. Guided by these admittedly anecdotal findings, we were able to find more convincing cases of instability. For example, the hit list “ONDER_CDH1_SIGNALING_VIA_CTNNB1” (Onder et al., 2008, 83 genes) has t_0_ vs t_now_ results with complete Jaccard similarity of 0.0, top-parents Jaccard of 0.024, and top-gene Jaccard of 0.056 - at the low extremes of all three. While the results are subjectively more dissimilar than the above examples (see Supplementary File 3), there are still prominent thematic similarities: the top terms at t0 include “angiogenesis”, “chemotaxis”, “locomotory behavior” and “skeletal development”, with replacements for some of these apparent at t_now_ including “blood vessel remodeling” and “cell chemotaxis”.

To summarize, as described in the main results section, by our objective measures many hit lists yield unstable enrichment results. But we find that on absolute scales, none of the measures adequately capture the subjective biological impression. Using a null was essential to calibrate the scores but was still not sufficient to address this disconnect. The “top term parents” measures had absolute scales that were more in line with the subjective impression hits lists give. But we feel there is room for improvement in developing semantic similarity measures and/or nulls that better capture the exploratory way in which GO enrichment is used, to the extent that the terms that are found are less important than the overall subjective impression. Because no measure seemed ideal for all possible applications, and some users may be more sensitive to differences in “impression” than us, we offer four measures on the GOTrack web site that can be plotted, and additional ways of plotting and exploring the data (see web site for details).

### Supplementary Figure legends

**Supplementary Figure 1:**
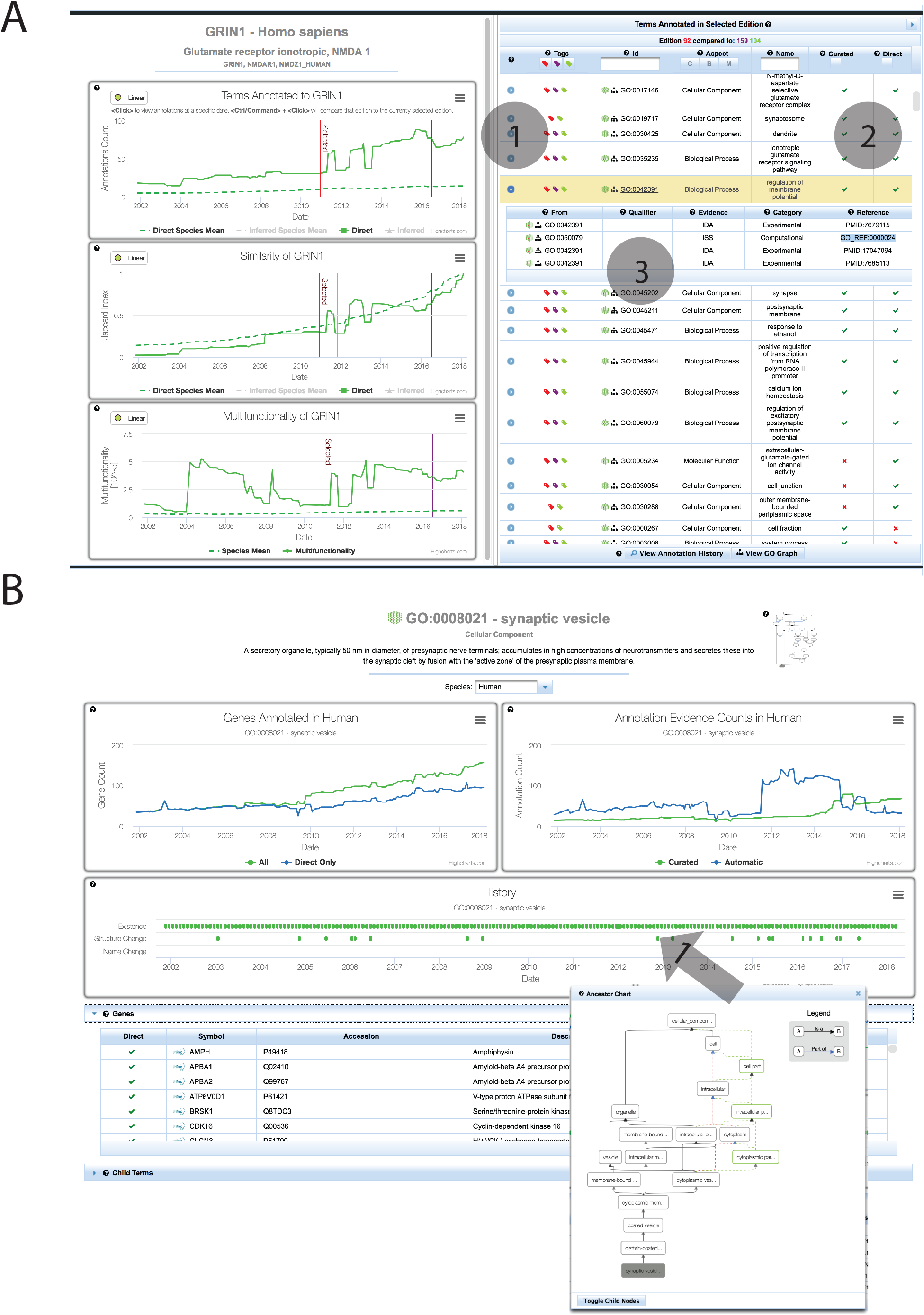
Screen shots of the gene and term tracking views of GOTrack. **A.** The main view of the Gene Tracker. At left are plots of annotation metrics over time. Clicking on a chart brings the focus to the selected time point. The table at right lists annotated GO terms. In this example, we are comparing three different time points, which are indicated as vertical lines on the plots, and colored tags in the table (1). The table indicates which annotations are direct and which are inferred (2). Expanding a row of the table (3) reveals details including evidence codes and annotation sources. **B.** The Term Tracker. The plots at the top of the page show number of genes annotated to the term (left) and number of annotations stratified by evidence type (right). The History timeline (centre) shows when the term was created in GO, and when changes were made. Clicking on a change point (Arrow 1) brings up the GO graph for the term with changes indicated in red or green.

**Supplementary Figure 2:**
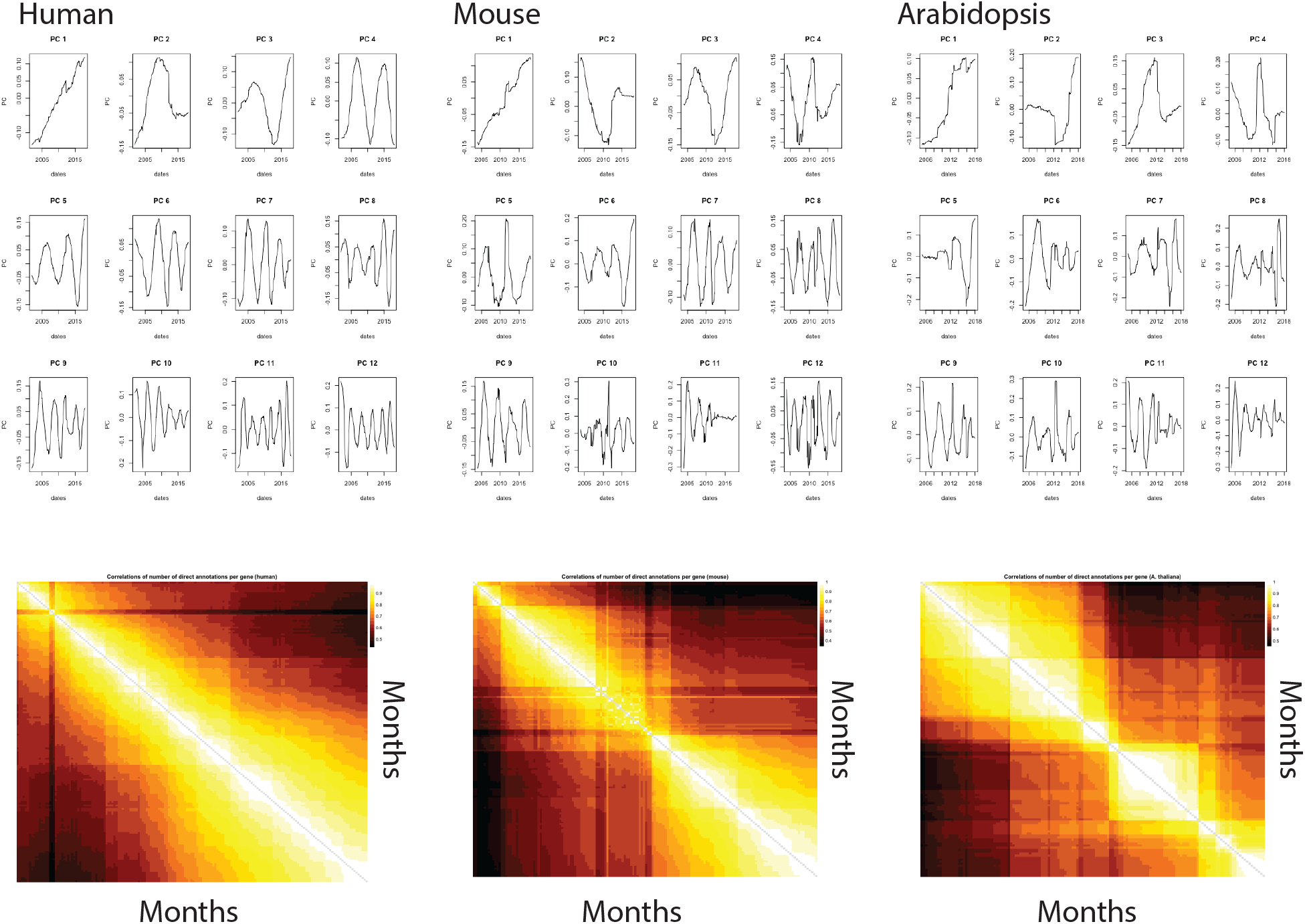
Correlation structure of direct annotation counts. Data are shown for three taxa. The **top** panel shows the profiles of the first twelve principal components of the direct annotation count matrix. PC1 reflects the general increase in annotations, while other components show more subtle periodic and transient changes. The heatmaps at the **bottom** are of the matrix of correlations of the direct annotation count vectors for each time point.

**Supplementary Figure 3:**
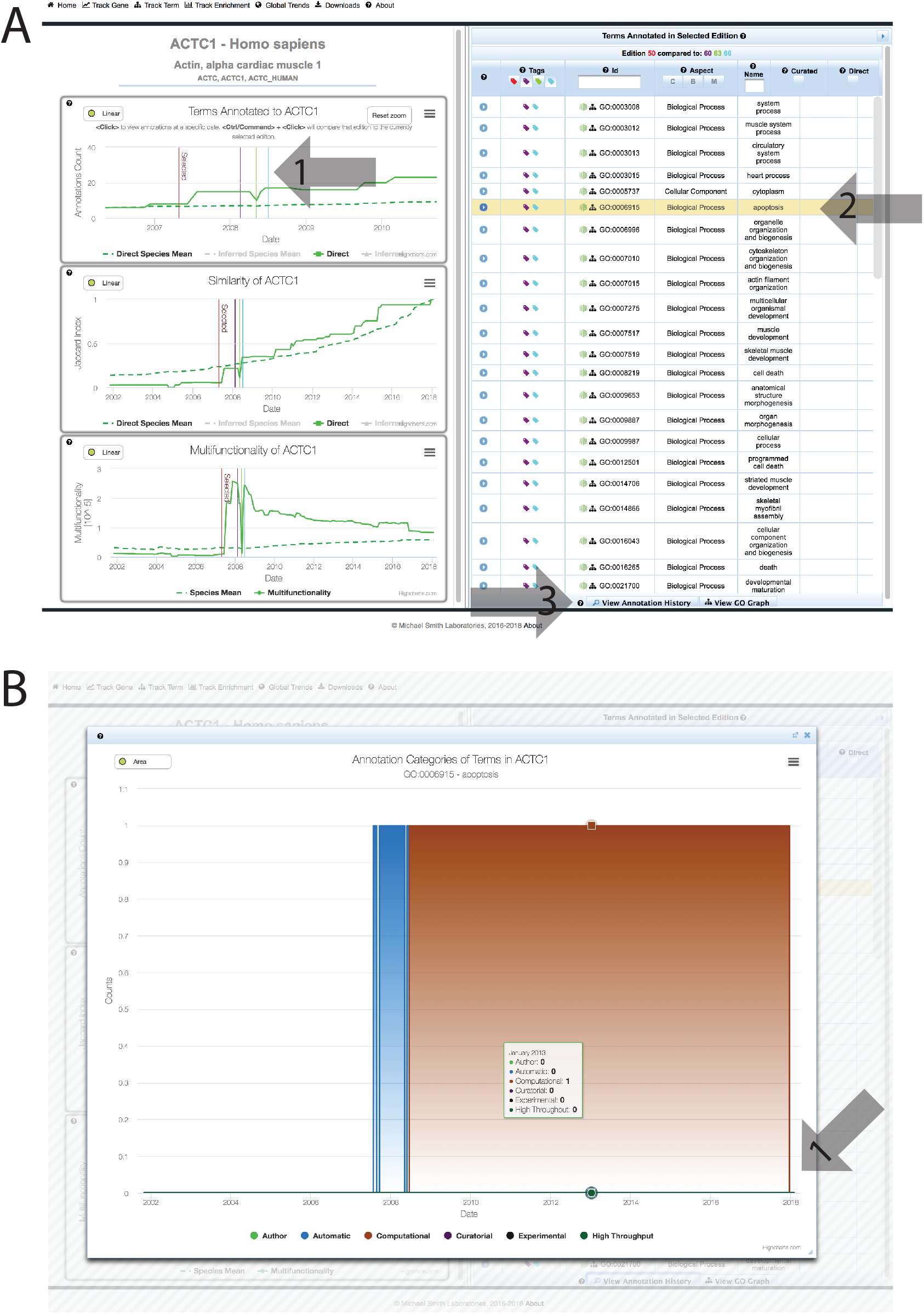
Exploring annotation volatility with GOTrack. **A.** The Gene tracker view for ACTC1, showing the comparison of multiple time points (vertical lines indicated by arrow 1). The term “apoptosis” was present in only two of the time points (Arrow 2), with the term missing from the middle time point. Clicking on the“View Annotation History” button (Arrow 3) brings up the history pane shown in B. **B.** The annotation history of “apoptosis” to ACTC1. The term was annotated to the gene but removed several times before the most recent removal in late 2017 (Arrow 1).

**Supplementary Figure 4:**
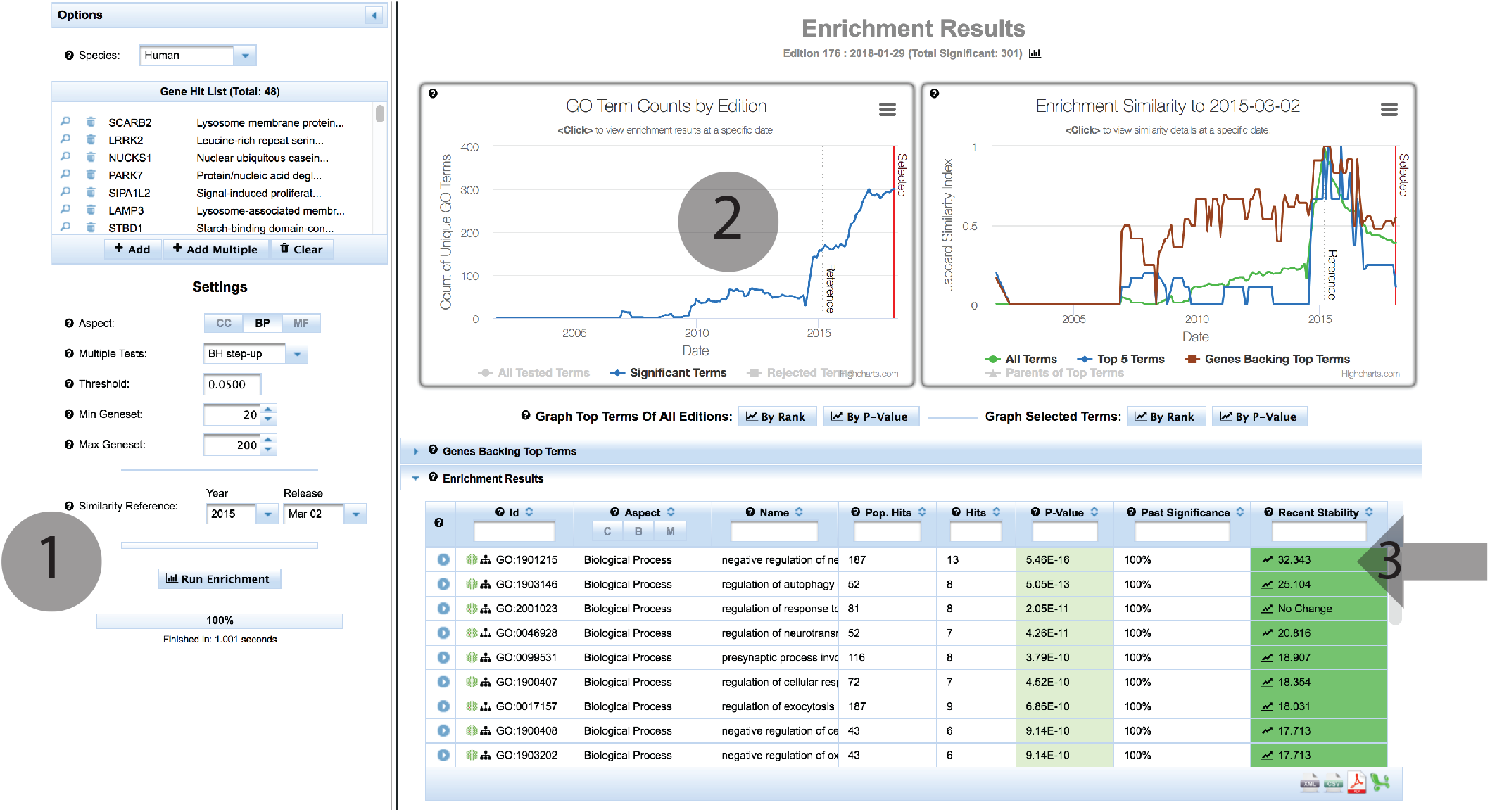
Screen shot of the GOTrack Enrichment Tracker. Users enter their genes and set parameters in the panel at left, including which date to use as a reference (1). Results appear at the right, with charts showing the number of significant terms (2) and similarity scores (upper right). Term-level information is available in the table (arrow 3).

**Supplementary Figure 5:**
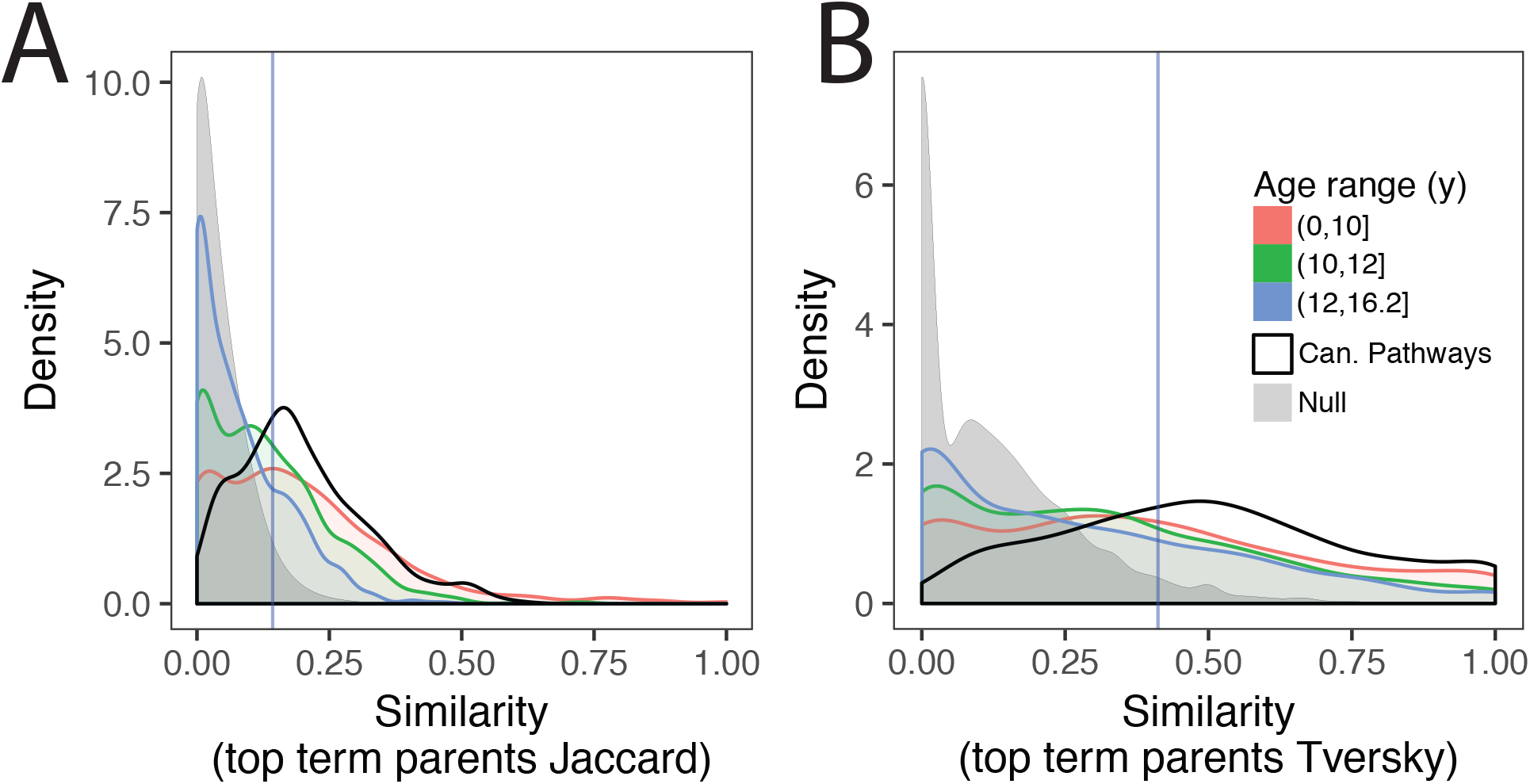
Stability analysis of the MSigDB hit lists using two additional measures of stability. Compare to Figure 4. **A:** The top-term parents Jaccard measure. **B.** The top-term parents Tversky measure. Vertical blue line indicates 95%ile of the null.

**Supplementary Figure 6:**
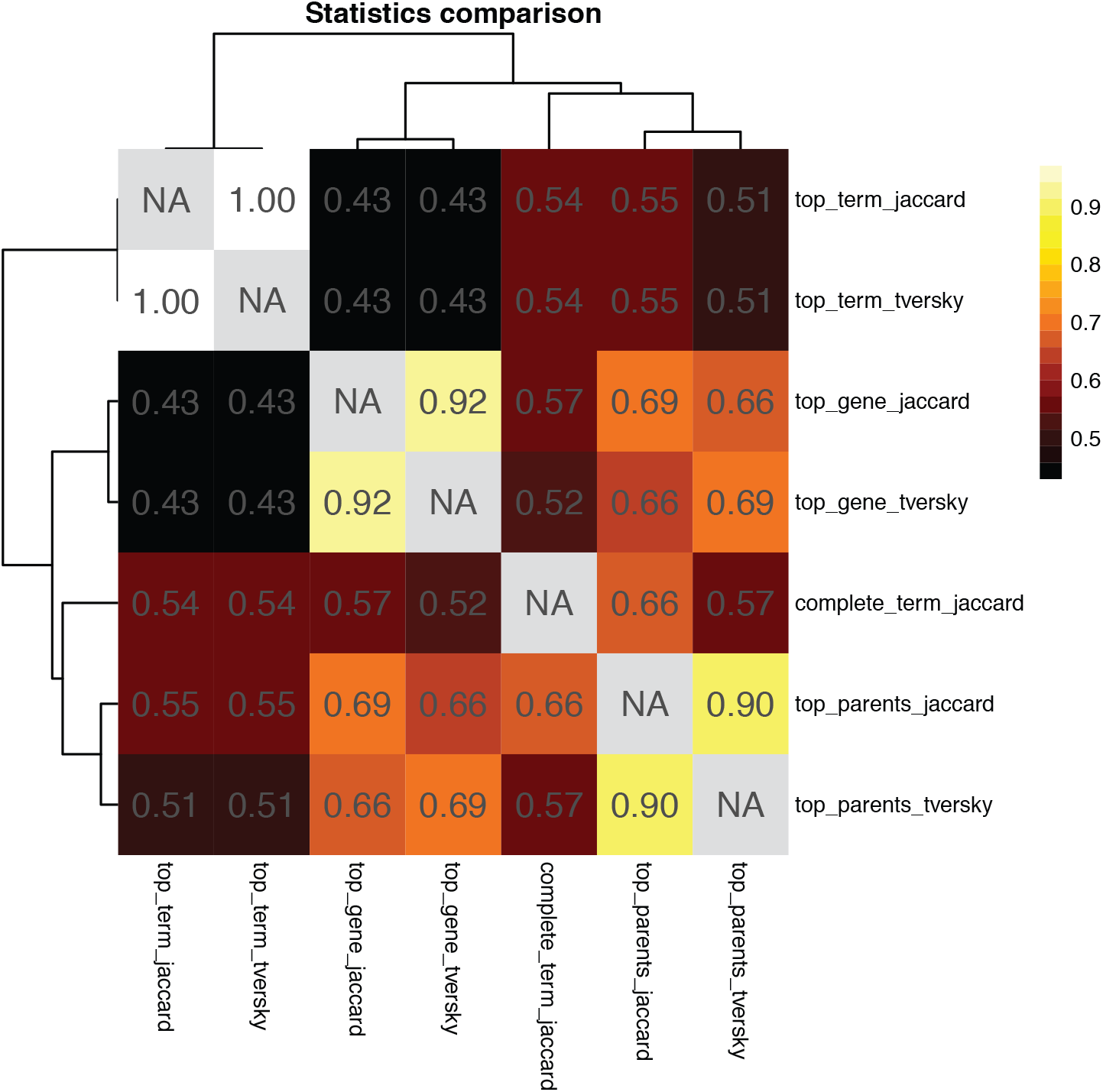
Summary of correlations among measures investigated, for the CGP hit lists.

### Supplementary File descriptions

Supplementary Files 1-3 (APPEL_IMATINIB_RESPONSE.enrichment.xlsx, BENPORATH_ES_2.enrichment.xlsx, ONDER_CDH1_SIGNALING_VIA_CTNNB1.enrichment.xlsx): Examples of CGP enrichment results discussed in the supplement. These files were downloaded from the GOTrack enrichment tracking tool, with highlighting added. Results for t_0_ and t_now_ are on separate tabs. Descriptions of the fields:

- ID, Aspect, Name: basic descriptors of the GO term.
- Term size: Number of genes annotated to the term in the analyzed edition.
- Hits annotated: Number of genes on the hit list annotated to the term in the analyzed edition.
- P-value: Enrichment p-value (uncorrected).

